# Illumination of the complement receptors C3aR and C5aR signaling by anaphylatoxins

**DOI:** 10.1101/2023.01.18.524551

**Authors:** Yue Wang, Weiyi Liu, Youwei Xu, Qingning Yuan, Xinheng He, Ping Luo, Wenjia Fan, Jinpeng Zhu, Xinyue Zhang, Xi Cheng, Yi Jiang, H. Eric Xu, Youwen Zhuang

**Affiliations:** The CAS Key Laboratory of Receptor Research, Center for Structure and Function of Drug Targets, Shanghai Institute of Materia Medica, Chinese Academy of Sciences, Shanghai 201203, China; University of Chinese Academy of Sciences, Beijing 100049, China; School of Chinese Materia Medica, Nanjing University of Chinese Medicine, Nanjing 210046, China; Lingang Laboratory, Shanghai 200031, China; School of Life Science and Technology, ShanghaiTech University, Shanghai 201210, China

## Abstract

The complement receptors C3aR and C5aR, whose signaling are selectively activated by anaphylatoxins C3a and C5a, are important regulators of both innate and adaptive immune responses. Dysregulations of C3aR and C5aR signaling lead to multiple inflammatory disorders, including sepsis, asthma, and acute respiratory distress syndrome (ARDS). The mechanism underlying endogenous anaphylatoxin recognition and activation of C3aR and C5aR remains elusive. Here we reported the structures of C3a-bound C3aR and C5a-bound C5aR1 as well as an apo C3aR structure. These structures, combined with mutagenesis analysis, reveal a conserved recognition pattern of anaphylatoxins to the complement receptors that is different from chemokine receptors, unique pocket topologies of C3aR and C5aR1 that mediate ligand selectivity, and a common mechanism of receptor activation. These results provide crucial insights into the molecular understandings of C3aR and C5aR1 signaling and structural templates for rational drug design for treating inflammation disorders.

---

The complement system represents a major part of innate immunity that plays critical role in host defense through cooperating with phagocytes and fluid antibodies to recognize and remove invading pathogens or damaged tissues, thus restoring the body homeostasis ^1–3^. In addition to innate response in vertebrates, the complement system also participates in opsonization and enhancement of adaptive immunity ^4,5^. Activation of the complement proteolytic cleavage cascades not only forms the membrane-attack complex (MAC) to directly kill pathogenic microorganisms, but also generates bioactive basic peptides with pro-inflammatory properties, including C3a and C5a, known as anaphylatoxins ^6,7^. Production of anaphylatoxins stimulate the activation of immune cells such as mast cells and basophilic leukocytes to release inflammation agents, like cytokines, chemokines, and histamine, which promote inflammation development. Anaphylatoxins also act as potent chemo-attractants for the migration of macrophages and neutrophils to the inflamed tissues, resulting in neutralization of the inflammatory triggers by multiple ways, such as phagocytosis and generation of reactive oxidants ^6,8–10^. The anaphylatoxins-induced local inflammation is essential for control of infection, nevertheless, upregulation of anaphylatoxin signaling also contributes to the development of many inflammatory disorders and autoimmune diseases, involving sepsis, acute respiratory distress syndrome (ARDS), allergic asthma, systemic lupus erythematosus (SLE), rheumatoid arthritis (RA), Alzheimer’s disease and even cancers ^6,8,11–13^. Regulation of anaphylatoxin signaling offer great potential in treating these pathological disorders.

C3a and C5a are two anaphylatoxin polypeptides with 77 and 74 amino acids, respectively, and they share 36% sequence similarity. Both C3a and C5a adopt a fold of four α-helices stabilized by three pairs of disulfide bonds ^14,15^. Despite the similarities in sequence and structure, C3a and C5a exert their functions by specifically binding to and activating different complement receptors, C3aR and C5aR, which belong to class A G protein-coupled receptors (GPCRs) ^16–19^. C5aR includes two subtypes, C5aR1 (CD88) and C5aR2 (C5L2 or GPR77), both show nanomolar affinity to C5a ^6,17,18^. As typical GPCRs, C3aR and C5aR1 primarily signal through G_i/o_ and conduct both G protein and arrestin signaling ^20–22^.

Due to the fundamental role of C3a-C3aR and C5a-C5aR signaling axes in innate and adaptive immunities, intensive efforts have been made to explore the binding and signaling properties of C3a and C5a to their respective receptors. Previous studies indicated that both C3a and C5a engaged the receptors by a “two-site” binding paradigm, involving the interactions with the transmembrane bundles and the extracellular regions, mainly including ECL2 ^23–25^. It was suggested that the C-terminus of C3a and C5a, especially the C-terminal arginine, are critical for receptor activation ^26,27^. Moreover, the N-terminal loop of C5aR1, but not that of C3aR, was necessary for efficient recognition with their respective anaphylatoxin ligands ^25,28^. Significant progresses were also achieved in structural studies on C5aR1, including crystal structures of C5aR1 bound to peptidomimetic antagonist PMX53 and allosteric antagonists avacopan and NDT9513727 ^29,30^. However, no structure of C3aR has been reported so far. The structural mechanisms of how C3a and C5a specifically bind and activate the complement receptors and downstream signaling transducer have long been sought after without success, which largely imped our understanding about anaphylatoxin-complement receptor signal axis and drug development targeting these important biological processes. Here, we reported cryo-EM structures of C3aR-G_i_ and C5aR1-G_i_ complexes bound to C3a and C5a, respectively, as well as the structure of C3aR-G_i_ complex in its apo state. These structures revealed the unique binding modes of C3a and C5a to their respective complement receptors, and the potential activation mechanism of C3aR and C5aR1, which may facilitate the rational design of anti-inflammation drugs targeting these receptors.

## Structure determination of C3aR-G_i_ and C5aR1-G_i_ complexes

To obtain the anaphylatoxin ligands C3a and C5a for assembling signaling complexes, we recombinantly expressed C3a and C5a in Sf9 insect cells and purified the proteins by Ni-NTA affinity chromatography. In this study, C3a and C5a were prepared as fusion proteins with a SUMOstar tag inserted onto the N terminus to facilitate expression and proper folding (Extended Data Fig. 1a,b) ^31,32^. To confirm whether the recombinant C3a and C5a are biologically functional, we performed cyclic AMP (cAMP) inhibition assays to test their capabilities in activating G_i_ signaling through C3aR and C5aR1 (Extended Data Fig. 1c). In our assays, both C3a and C5a could suppress cAMP accumulation in dose-dependent manners through C3aR and C5aR1 with EC50 values at 1.0 nM and 0.1 nM, respectively (Extended Data Fig. 1c). While C3a and C5a showed over 1000-fold selectivity to their respective receptors, C3a activated C5aR1 with relative lower efficacy than C5a (~30% of Emax of C5a) (Fig. 1b). Our results are consistent with previous investigation on the selective activities of C3a and C5a ^28,33^. We then assembled the C3aR-G_i_ and C5aR1-G_i_ complexes in membrane by co-expressing the receptor with G_i_ heterotrimer and determined the structures by cryo-EM (Fig. 1c-1e and Extended Data Fig. 1d-f). The single chain antibody scFv16 was added to stabilize the whole complexes. The structures of apo state C3aR-G_i_, C3a-bound C3aR-G_i_, and C5a-bound C5aR1-G_i_ complexes were determined at the resolution of 3.2 Å, 2.9 Å, and 3.0 Å, individually (Extended Data Fig. 2). The density maps allowed clear definition and modeling of most portions of the receptors, the anaphylatoxin ligands, scFv16 antibody, and G_i_ heterotrimer without the flexible alpha-helix domain of G_αi_ (Extended Data Fig. 3). In the structures, both C3a and C5a comprised a core of four α helices stabilized by three disulfide bonds, agreed well with their crystal structures ^14,34^ (Extended Data Fig. 4a, b). Compared to other anaphylatoxin receptors, C3aR contains an extra-large second extracellular loop (ECL2, about 172 amino acids), in which only the first 16 residues of ECL2 (V159 to K175) were resolved, indicating the extremely dynamic property of ECL2 in C3aR, possibly due to the lack of interaction with C3a ligand.

**Fig. 1.**
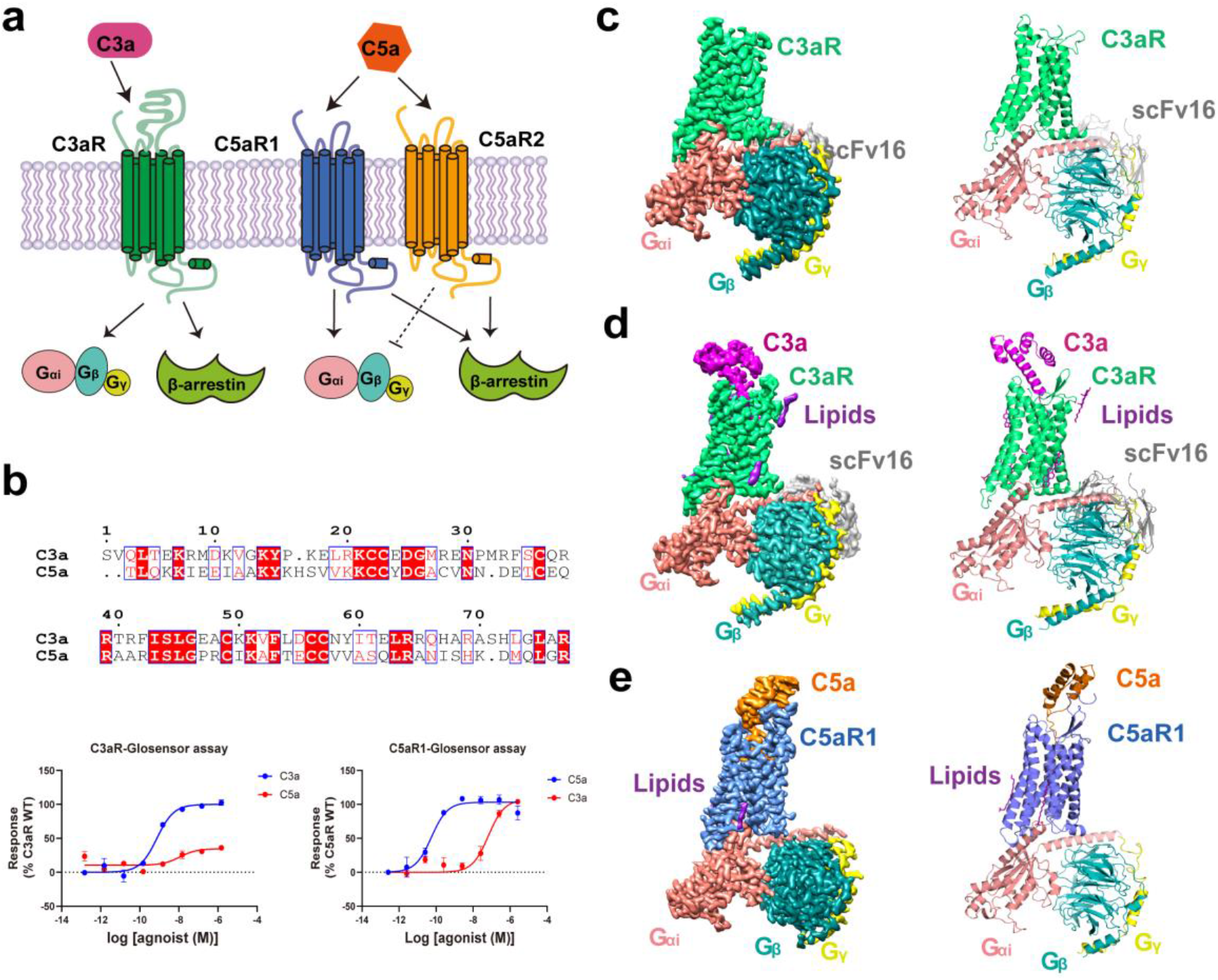
Overall structures of the C3aR-G_i_ and C5aR1-G_i_ complexes. **a,** Cartoon presentation of C3aR and C5aR signaling pathway mediated by C3a and C5a. **b,** Activities of C3aR and C5aR1 induced G_i_ signaling by C3a and C5a. Upper panel: sequence alignment of human C3a and C5a. Lower panel: the dose-dependent response curves of C3aR and C5aR1 activated by C3a and C5a using GloSensor cAMP assays. Data shown are means ± S.E.M. from three independent experiments performed in technical duplicate. **c, d, e,** Orthogonal views of the cryo-EM maps (left panels) and models (right panels) of the apo-C3aR-G_i_ complex (**c**), C3a-C3aR-G_i_ complex (**d**) and C5a-C5aR1-G_i_ complex (**e**).

## Recognition of C3a by C3aR

C3a recognizes and activates C3aR with nanomolar affinity ^35^. The whole C3a molecule occupied an amphipathic pocket with a size of 1297 Å^3^, which is composed by residues from TM2, TM3, TM5-7, ECL2 and ECL3 from the extracellular half of C3aR. It was shown that the C terminal pentapeptide ‘LGLAR’ contains the active-site of C3a, which is indispensable for C3aR activation ^36,37^. Consistently, in our structure, this pentapeptide of C3a occupied the orthosteric pocket in the transmembrane domain (TMD) in a C-terminus-inside mode. Binding of C3a to C3aR is mainly mediated by two independent regions, namely C3aS1 and C3aS2 (Fig. 2a), which together contributes to the high potency of C3a to C3aR.

**Fig. 2.**
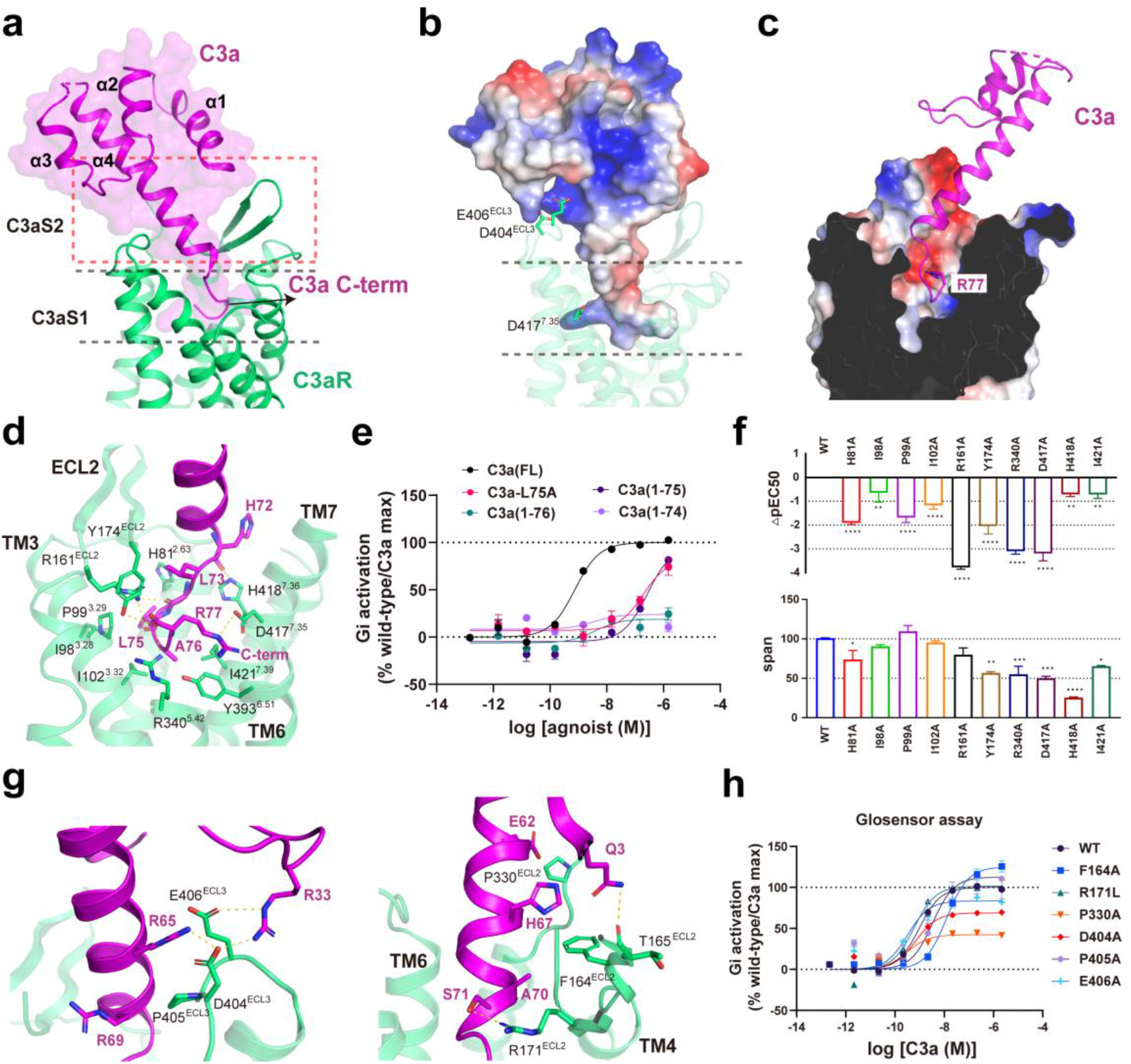
Recognition of C3a by C3aR. **a,** Two-site binding regions of C3a in C3aR. **b,** Electrostatic surface representation of C3a. **c,** Cross-section of C3a binding pocket, R77 of C3a is shown as stick. **d,** Interactions between “HLGLAR” of C3a and C3aS1 of C3aR. **e,** The dose-dependent response curves of C3aR induced by different length of C3a measured by GloSensor cAMP assay. Data shown are means ± S.E.M. from three independent experiments performed in technical duplicate. **f,** Effects of mutants in the C3aS1 induced by C3a on cAMP response. Data are presented as means ± S.E.M. of three independent experiments performed in technical duplicate. All data were analyzed by two-side, one-way ANOVA by Fisher’s LSD test compared with WT. *P<0.05; **P<0.01 and ***P<0.001 were considered as statistically significant. **g,** Interaction of the C3a core region with C3aR. Left panel: the polar interactions between C3a and ECL3 of C3aR. Right panel: interactions between C3a and ECL2 of C3aR. Yellow dash indicates the hydrogen bonds. **h,** The dose-dependent response curves of mutants in C3aS2 of C3aR induced by C3a measured by GloSensor cAMP assay. Data shown are means ± S.E.M. from three independent experiments performed in technical duplicate.

In C3aS1, the terminal residues ‘LGLAR’ adopts a ‘hook’ conformation with the last arginine residue pointing toward a negatively charged sub-pocket containing D417^7.35^ (Fig. 2b) (receptor residues are noted with superscript based on Ballesteros-Weinstein numbering method ^38^ and residues of C3a and C5a are noted without superscript), fixing the C3a molecule into the receptor TMD core. In addition to ionic interactions with D417^7.35^, the side chain of R77 also forms *cation-π* interaction with Y393^6.51^ while its main chain carboxylate forms hydrogen bonds with the side chains of Y174^ECL2^ and R340^5.42^ (Fig. 2c, d). Mutations of D417^7.35^, Y174^ECL2^, R340^5.42^ and Y393^6.51^ to alanine or deletion of R77 from C3a greatly decreased G_i_ activation by C3aR (Fig. 2e, f and Supplementary Table 2). The extensive interactions of R77 with C3aR are consistent well with previous results, which indicated that R77 is a critical determinant for C3aR activation ^26^. The main chain of H72, G74 and L75 also formed hydrogen-bonds with H418^7.36^ and R161^ECL2^, respectively (Fig. 2d). Consistently, mutation of H418^7.36^A diminished the efficacy of C3a while R161^ECL2^A decreased the potency of C3a to C3aR by over 100 folds (Fig. 2f). In addition to polar interactions, L73, L75 and A76 in C-terminal hook of C3a made hydrophobic contacts with nearby residues I98^3.28^, P99^3.29^, I102^3.32^, Y174^ECL2^ and I421^7.39^ within the orthosteric binding pocket (OBP) (Fig. 2d, f), and mutations in these pocket residues also resulted in decrease of C3a-mediated C3aR activation (Fig. 2f, h and Supplementary Table 2).

Engagement of C3a with C3aR was further enhanced by binding in C3aS2, which located in the extracellular vestibular of C3aR (Fig. 2a). The C3aS2 comprised interaction between ECL2 and α4 helix of C3a, as well as interactions of ECL3 with the α4 helix and α3-α4 loop of C3a. It was worth noting that D404^ECL3^ and E406^ECL3^ inserted into a positively charged cavity near α4 helix and α3-α4 loop regions of C3a, forming close salt bridges with R33 and R65 (Fig. 2g). Nevertheless, the mutagenesis data showed that mutations of C3aS2 residues had less significant effects on C3a activity compared to those in C3aS1 (Fig. 2h), consistent with the relatively poor resolved density of C3a in the region outside of the C-terminal hook and α4 helix.

## Conservation and divergency in C3a and C5a binding modes

C5a shared the conserved C-terminal pentapeptide sequence with C3a (Fig. 1b), implying the potential conserved molecular patterns in recognition of C3a and C5a to their receptors. Likewise, in our structure, C5a also bound to the cognate receptor C5aR1 in C-terminus-inside mode (Fig. 3a). All the residues of the C-terminal loop contributed to the potent recognition of C3a and C5a to their cognate receptors. Computational simulations indicated that both C3a and C5a stably bound to the OBPs of their cognate receptors, with RMSD values of the poses in cryo-EM structures ranging from 1 Å to 2 Å (Extended Data Fig. 4c). The C-terminal pentapeptide ‘MQLGR’ of C5a occupied a ‘hook’ shaped configuration, similar to ‘LGLAR’ of C3a in C3aR. Structural alignment revealed that C3a and C5a overlaid well in the C-terminal hook (Fig. 3b), with R74 of C5a engaging a conserved negatively charged sub-pocket in C5aR1 as that occupied by R77 of C3a in C3aR. Despite the similarities, remarkable differences were observed in C5a binding mode. Compared to C3a in C3aR, the C-terminal loop of C5a was longer with five additional residues (decapeptide), which is stretched and formed broader interactions with the ligand pocket of C5aR1 (Fig. 3c). Additionally, unlike C3aR, the N-terminal loop of C5aR1 made direct interactions with C5a, consequently, the α-helix core of C5a adopted a clockwise rotation of ~84° toward the N-terminal loop of C5aR1 (as measured by the relative conformation of α4 helix of C3a and C5a) (Fig. 3b).

**Fig. 3.**
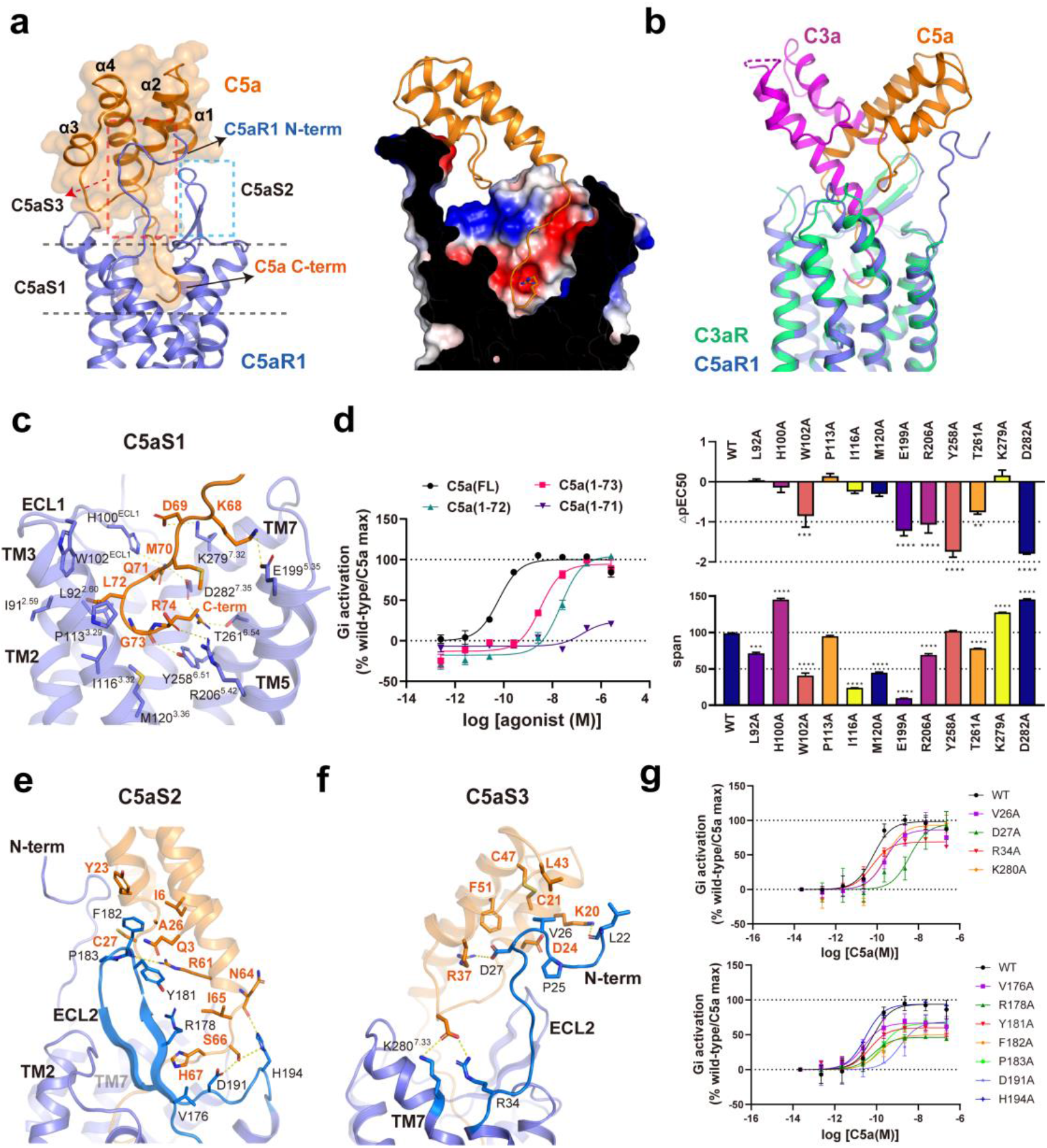
Recognition of C5a by C5aR1. **a,** Interactions between C5a and C5aR1. Left panel: three-site binding regions of C5a in C5aR1. Right panel: cross-section of C5a binding pocket, R74 of C5a is shown as stick. **b,** Structural superposition of C3a-bound C3aR and C5a-bound C5aR1. **c,** Interactions between C-terminal loop “KDMQLGR” segment of C5a and C5aS1 of C5aR1. **d,** Left panel: effects of C-terminal residue truncation of C5a on downstream signaling of C5aR1. Right panel: Effects of mutants in the C5aS1 induced by C5a on cAMP response. Data are presented as means ± S.E.M. of three independent experiments performed in technical duplicate. All data were analyzed by two-side, one-way ANOVA by Fisher’s LSD test compared with WT. *P<0.05; **P<0.01 and ***P<0.001 were considered as statistically significant. **e,** Interactions between C5a with C5aS2 of C5aR1. **f,** Interactions between C5a with C5aS3 of C5aR1. **g,** The dose-dependent response curves of mutants in C5aS2 (upper panel) of C5aR1 and mutants in C5aS3 (lower panel) of C5aR1 induced by C5a measured by GloSensor cAMP assay. Data shown are means ± S.E.M. from three independent experiments performed in technical duplicate.

EM density map was clearly defined across the whole C5a molecule in the C5a-C5aR1 complex structure. Compared to C3a in C3aR, C5a possessed in a much wider amphipathic pocket in C5aR1 with a size of 1744 Å^3^, consistent with 10-fold higher potency of C5a than C3a to their cognate receptors (Fig. 1b and Extended Data Fig. 1c). The interaction interface of C5a in C5aR1 is consist of three separated sites: namely C5aS1, C5aS2 and C5aS3, among which C5aS1 constituted the major binding site for C5a (Fig. 3a). All the three binding sites appeared to be amphipathic. The C-terminal octapeptide of C5a, which was identified as the smallest fragment for reasonable C5aR1 binding ^39^, occupied C5aS1 site and interacted with residues from TM2, TM3, TM5-7 and ECL1 of C5aR1 (Fig. 3c). In C5aS1, conformation of the C-terminal octapeptide loop of C5a was stabilized by a widespread polar interaction network, including the ionic patches formed by K68, D69 and R74 with residues E199^5.35^, K279^7.32^, R206^5.42^ and D282^7.35^, respectively, as well as a hydrogen bond network formed by Q71, G73 and R74 with residues H100^ECL1^, Y258^6.51^, T261^6.54^, and D282^7.35^ (Fig. 3c). Consistent with the binding pose in our structure, mutations of individual residues in the polar network all showed compromised activity in G_i_ activation, especially Y258^6.51^ and D282^7.35^ (Fig. 3d). To further verify the structural findings, we generated residue-by-residue C-terminal truncations of C5a from R74 to M70, and tested their effects on G_i_ signaling of C5aR1 (Fig. 3d). The data suggested that these C5a truncations exhibited linearly decreased activities in inducing C5aR1 activation and truncation at L72 nearly eliminated C5aR1 signaling, supporting the binding mode of C5a in C5aS1 (Fig. 3d). Our structural and mutagenesis data are consistent with the previous result that the C-terminus of C5a, particularly R74, was indispensable for effective C5aR1 activation by C5a ^24,40^, and mutations on E199^5.35^, R206^5.42^ and D282^7.35^ significantly reduced C5a binding affinity to C5aR1 ^41–45^. In addition to the extensive polar contacts, the side chain of L72 extended toward the cleft of TM2 and TM3, and inserted into a hydrophobic pocket formed by residues I91^2.59^, L92^2.60^, W102^ECL1^, P113^3.29^ and I116^3.32^ (Fig. 3c).

In our structure, clear densities of the complete ECL2 and the N-terminal loop from L22^N-terminus^ to R34 ^N-terminus^ of C5aR1 were resolved, constituting the C5aS2 and C5aS3 sites for C5a binding, respectively. The ECL2 of C5aR1 interacted broadly with residues from α1, α2, α2-α3 loop, α4 and part of C-terminal loop of C5a while the N-terminal loop interacted with residues from α2-α4, α2-α3 loop and α3-α4 loop of C5a (Fig. 3a). The N-terminal loop of C5aR1 was previously suggested to be the second site for efficient C5a binding in addition to the TMD pocket engaged by C terminus of C5a ^24,25,46–50^. In the structure, D27 ^N-terminus^ and R34 ^N-terminus^ of C5aR1 made salt bridge interactions with R37 in α3 helix and D31 in α2-α3 loop of C5a, respectively (Fig. 3f). Mutation of D27 ^N-terminus^ A and deletion of the N-terminal 33 or 34 residues in C5aR1 largely diminished or abolished the G_i_ activation by C5a (Fig. 3g and Supplementary Table 3), suggesting the indispensable role of D27 ^N-terminus^ in C5a binding, which was in accordance with the structure as well as the previous mutagenesis and NMR studies toward N-terminus of C5aR1 ^47–49^. However, no obvious decrease of C5a activity was observed in R34 ^N-terminus^ A mutant of C5aR1, indicating the limited influence of this residue in C5aR1 activation by C5a (Fig. 3g). Apart from C5aS2, C5aS3 provided another independent anchoring site for C5a binding and enrichment. In C5aS3, the main chain carbonyl of F182^ECL2^ with the side chains of D191^ECL2^ and H194^ECL2^ formed hydrogen-bond interactions with R61, N64 and S66 of C5a (Fig. 3e). Furthermore, F182^ECL2^ and P183^ECL2^ formed extensive hydrophobic interactions with nearby C5a residues, including Q3, I6, Y23 and C27 (Fig. 3e). Accordingly, mutations of the above residues in C5aS3 to alanine decreased C5a activity (Fig. 3g).

## Distinct binding modes of C3a and C5a from chemokines

Similar to chemokines, C3a and C5a belong to large macromolecular ligands of GPCRs and behave as strong chemotaxis for immune cells. The molecular recognition of chemokines to their cognate chemokine receptors were previously investigated ^51–56^. Whereas recognitions of chemokines by chemokine receptors exhibit diversities in binding modes such as differences in the depth of the chemokine N-termini into the TMD pockets, they share the conserved “two site” model, with the N-terminal loop of the chemokine insert into the orthosteric TMD pocket while the globular core region interacts with the N-terminal segments of the chemokine receptors. In contrast to the N-terminus-inside mode of chemokines to their receptors, both C3a and C5a recognize the TMD pockets of their receptors in the C-terminus-inside mode. Structural comparison of C3a-C3aR or C5a-C5aR1 with chemokine-bound receptors revealed that the C-terminal loops of C3a and C5a insert into the orthosteric TMD pockets, with similar depth as the N-termini of CCL2, CCL3 and CCL15 into their cognate chemokine receptors ^54–56^ (Extended Data Fig. 5a, b). In our structures, the last C-terminal arginine of C3a and C5a occupied a negatively charged sub-pocket in the bottom of the OBP in part composed by D^7.35^, which is only conserved among anaphylatoxin receptors but not in chemokine receptors. Correspondingly, the first N-terminal residues of chemokines are not conserved and fit into an amphipathic or a hydrophobic sub-pocket in the bottom of OBPs of chemokine receptors (Extended Data Fig. 5c).

## Ligand induced activation mechanisms of C3aR and C5aR1

The crystal structure of C5aR1 bound to peptidomimetic antagonist PMX53 was previously reported ^29,30^. Structural alignment of C5a and PMX53 bound C5aR1 structures revealed important conformational changes of C5aR1 activation induced by C5a binding. Compared to PMX53, C5a bound ~ 6Å deeper into the OBP of C5aR1 when measured at the Cα of the C-terminal arginine (Fig. 4a, b). As a result, the side chain of Y393^6.51^ of C5aR1 adopted an obviously downward shift related to the PMX53 bound state, facilitated by the close *cation-π* interaction with R74 of C5a (Fig. 4b). Conformational change of Y393^6.51^ was accompanied by the downward rotated movement of the toggle switch residue W390^6.48^ and F251^6.44^ of the P^5.50^I^3.40^F^6.44^ core triad (Fig. 4b and Extended Data Fig. 6a). C5a binding also induced twist and outward movement of TM3, mainly due to the outwardly stretched hydrophobic side chain of L72 of C5a into the cleft of TM2 and TM3 (Fig. 4c), which is accompanied by an inward movement of TM7 (Extended Data Fig. 6c). The above conformational changes subsequently caused rearrangement of the P^5.50^I^3.40^F^6.44^ triad, the outward kink of intracellular region of TM6, the collapse of Na^+^ pocket and the alteration of DR^3.50^F motif and conserved NPxxY motif (Fig. 4c and Extended Data Fig. 6), which ultimately lead to the opening of the intracellular cavity to accommodate G protein (Extended Data Fig. 6g). Helix 8 from the inactive C5aR1 structure was shown to maintain a reversed orientation toward the intracellular center of the TM bundles, restricting the engagement of G protein to C5aR1. In our structure, we observed that the side chain of Y300^7.53^ adopted a down shifted conformation, which rearranges helix 8 out of C5aR1 intracellular center (Extended Data Fig. 6h), thus releasing the space in intracellular cavity for G protein coupling. Conformation of Y300^7.53^ was stabilized by hydrophobic packing with F75^2.43^ and hydrogen bond interaction with R310^C-terminus^ (Extended Data Fig. 6c).

**Fig. 4.**
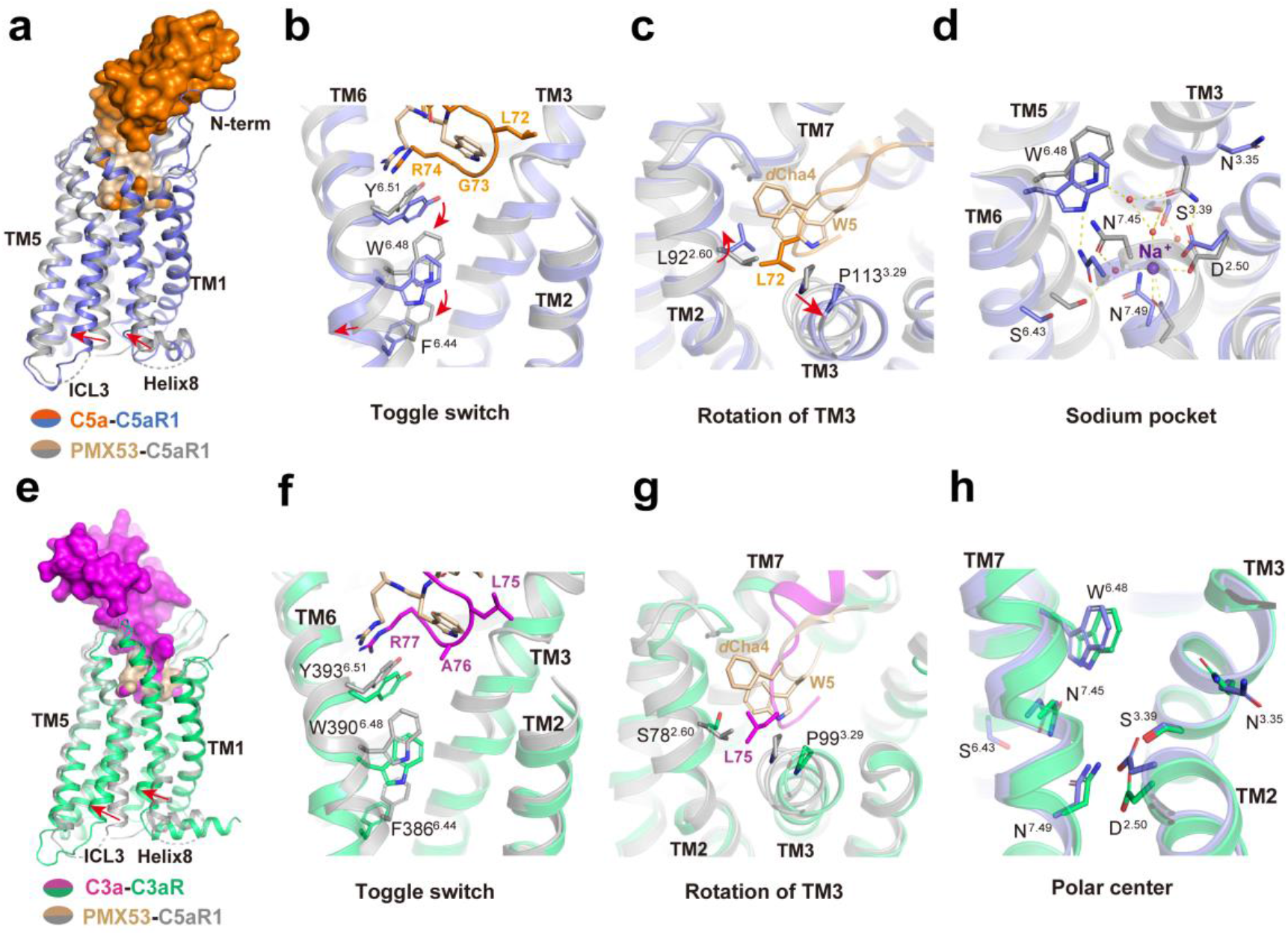
Ligand induced activation mechanisms of C3aR and C5aR1. **a,** Structural superposition of C5a-bound C5aR1 and PMX53-bound C5aR1 (PDB: 6C1R). **b,c,d,** Conformational changes upon C5aR1 activated by C5a. **(b)** Toggle switch; **(c)** rotation of TM3; **(d)** collapse of sodium pocket. The conformational changes of residues are shown as red arrows upon receptor activation. **e,** Structural superposition of C3a-bound C3aR and PMX53-bound C5aR1 (PDB: 6C1R). **f, g,** Conformational changes upon C3aR activated by C3a. **(f)** Toggle switch; **(g)** rotation of TM3. **h,** The similar polar center within the sodium pocket formed by activated C3aR and C5aR1.

Structure of C3aR shared high similarity with C5aR1, with RMSD of 0.763 Å when measured at the Cα atoms of the TMD region. Superposition of the C3a-bound active C3aR and PMX53-bound inactive C5aR1 structures revealed similar sets of conformational changes in C3aR activation as compared to those of C5aR1 (Fig. 4e), including the toggle switch and outward kink of TM6 (Fig. 4f), the variation of polar interaction network beneath the OBP, as well as the rearrangement of the PV(I)F motif and DRX motif (Extended Data Fig. 6d-f). Together, these conformational changes indicated that C3aR may share conserved activation mechanism as C5aR1 (Extended Data Fig. 6i). We also obtained a structure of apo C3aR coupled with G_i_ heterotrimer. Despite the noticeable differences in the extracellular vestibules due to C3a binding, structures of C3a-bound and apo C3aR highly resembled each other in the TMD core and intracellular part (Extended Data Fig. 8a). Structural superposition of C3aR in its active and apo states revealed the potential structural determinants responsible for the constitutive activity of C3aR. In the apo-C3aR structure, due to the absence of interaction with ligand, the polar side chain of R340^5.42^ shifted toward TM6 and formed close hydrogen bond interaction with Y393^6.51^, which is critical for the receptor activation (Extended Data Fig. 8d and Supplementary Table 2). The polar connection of R340^5.42^ and Y393^6.51^ stabilizes the conformation of Y393^6.51^ in its active state, which is consistent with the high level of constitutive activation of C3aR (Extended Data Fig. 8e).

## G_i_ coupling of C3aR and C5aR1

Structural alignment of C3aR-G_i_ and C5aR1-G_i_ complexes indicated that the G_i_ coupling interfaces of C3aR and C5aR1 quite resemble each other (Extended Data Fig. 9a). The overall conformation of G_i_ coupling to C3aR and C5aR1 were suggested to be in the canonical states as observed in most of the GPCR-G_i_ complex structures reported, in contrast to the non-canonical states observed in structures of NTSR1 or GHSR complexed with G_i_ heterotrimer ^57,58^. In spite of the conformational similarity, subtle structural difference could be seen in the G_i_ coupling interface, with the α5 of G_αi_ subunit inserting deeper into C3aR intracellular cavity and shifting toward TM6 at about 1.5Å compared to those in C5aR1 (Extended Data Fig. 9b). The structural difference in G_i_ was possibly due to the anaphylatoxin ligand induced subtle conformational variation of the extracellular regions of C3aR and C5aR1, specifically, a more outward movement at about 2Å of TM6 extracellular end of C3aR relative to C5aR1 (Extended Data Fig. 9c). The G_i_ binding interfaces of C3aR and C5aR1 mainly include interactions of residues from TM3, TM5, TM6, ICL2, and ICL3 of the receptors and residues in β2-β3 loop, β6, and α5 helix of G_αi_ subunit (Extended Data Fig. 9). The ICL3 of C3aR and C5aR1 showed clear divergencies in structural topologies and interaction mode with G_i_, with the ICL3 of C3aR adopted broader interaction with the β6 and α5 helix of G_αi_ (Extended Data Fig. 9f and 9i).

## Summary

Signal transductions mediated by anaphylatoxins C3a and C5a, and their receptors constitute an essential part of immune responses for generating local inflammation and resolving infection. The C3a-C5a signaling axis has long been served as important targets for multiple inflammation disorders. In this paper, we reported the relatively high-resolution structures of C3aR and C5aR1 either in the apo or in the C3a and C5a bound states. These structures provide insight into the unique C-terminus inside binding modes of C3a and C5a to their receptors, in contrast to the N-terminus inside mode occupied by other chemoattractant peptides, such as formyl-peptides^59–61^ and chemokines^51–56^. Like many chemokine-bound to their receptors, the N-terminal loop of C5aR1, rather than that of C3aR, serves as important anchoring site for high affinity binding to C5a. The structures, together with mutational data, also revealed the highly conserved recognition pattern of C3a and C5a in the C3aR and C5aR1 binding pockets, with the last C-terminal arginine residue occupying the bottom negatively charged pocket, which is formed by residues including conserved residues R^5.42^, Y^6.51^, and D^7.35^. Our results provide comprehensive structural basis of the pharmacology and signaling of complement receptors C3aR and C5aR1 signaling and multiple structural templates for rational drug development targeting the complement system.

## Materials and Methods

### Generation of recombinant C3a and C5a

The recombinant wild-type C3a, C5a and a series of C3a/C5a mutants were generated in similar methods. The coding sequence of human C3a (residues1-77) was modified with a glycoprotein 67 (gp67) signal peptide followed by His6-tag and SUMOstar-tag in the N-terminus. The coding sequence of human C5a (residues1-74) was cloned in the same strategy. C3a and C5a constructs were all cloned into pFastbac1 vector (ThermoFisher) and expressed in Sf9 insect cells as secreted proteins using baculovirus infection system. The media was harvested after infection at 48 hours. The pH of supernatant was balanced by adding 1M HEPES (pH 7.4). For quenching the chelates, 1 mM nickel chloride and 5 mM calcium chloride were added and stirred for 1 hour at 4°C. Resulting precipitates were removed by centrifugation at 8,000 rpm (JA-10) and the supernatant was loaded onto Ni-NTA and incubated overnight. The Nickel resin was washed with buffer containing 20 mM HEPES pH 7.4, 100 mM NaCl and 40mM imidazole for 10 column volumes and then eluted in the above buffer containing 300mM imidazole. The eluted C3a/C5a were concentrated and purified over a size exclusion chromatography using a Hiload 16/600 superdex 75pg column. C3a/C5a peak fractions were pooled, concentrated and fast-frozen by liquid nitrogen and stored at −80 °C for further usage.

### Preparation of apo/C3a-C3aR-G_i_ and C5a-C5aR1-G_i_ complex

The full length human C3aR (residues 1-482) and C5aR1 (residues 1-350) was used to obtain C3a and C5a bound G_i_ complex, respectively. For apo state of C3aR-G_i_ complex, the C3aR was truncated to residues 1-476. The N-terminus of both C3aR and C5aR1 were modified with prolactin precursor sequence as a signal peptide, followed by FLAG-tag and fragment of β_2_AR N-terminal tail region (BN, hereafter) as fusion protein to increase the protein expression. The C-terminus of C3aR and C5aR1were fused with His×8 tag. A dominant-negative bovine G_αi_1 (G_αi_1_2M) with two mutations (G203A and A326S ^62^) was generated by site-directed mutagenesis to decrease the affinity of nucleotide binding and limit G protein dissociation for stable receptor-Gi complex. The N-terminus of rat Gβ1 was fused with a His×8 tag for two-step purification. All of the components of G_i_ heterotrimer, G_αi_1_2M, His8-G_β1_ and bovine G_γ2_, were cloned into pFastbac1, respectively.

The single chain antibody scFv16 was applied to improve the stability of the protein complex through enhancing the interface between G_αi_1 and G_β1_. The scFv16 antibody was prepared based on the method as previously reported ^63^. Briefly, secreted scFv16 was purified from expression media of baculovirus-infected Sf9 insect cells culture using Ni-NTA and size exclusion chromatography. After removing the chelates by Ni^2+^ and Ca^2+^, the supernatant from 2-liter culture was collected and loaded onto a gravity column with 5mL Ni-NTA resin. The nickel resin was first washed with 20mM HEPES pH 7.2, 100mM NaCl, 50mM imidazole for 10 column volumes and then eluted in buffer containing 300mM imidazole. The elution was concentrated to 2mL using centrifugal filters with a 30 kDa molecular weight cut-off (ThermoFisher) and applied to a HiLoad Superdex 200, 10/60 column (GE Healthcare). The monomeric peak fractions were collected, concentrated and fast-frozen by liquid nitrogen as stocks for further usage.

For C5a-C5aR1-G_i_ complex, SUMOstar-C5a, C5aR1, G_αi_1_2M, His8-G_β1_ and G_γ2_ were co-expressed in Sf9 insect cells at a ratio of 1:1:1:1:1 when the cell density reached to 4×10^6^ cells/mL. For C3aR-G_i_ complex, C3aR, G_αi_1_2M, His8-G_β1_ and G_γ2_ were co-expressed for preparation of the apo or C3a bound C3aR-G_i_ complex in the same infection virus ratio and cell density as C5a-C5aR1-G_i_ complex. After infection at about 48 hr, the cells were collected by centrifugation at 2000 g (ThermoFisher, H12000) for 20 min and the pellets were stored at −80°C for further purification. In preparation and formation of the C3a-bound C3aR-G_i_ complex, the SUMOstar-C3a was added during the protein purification.

For the purification of C3a-C3aR-G_i_ complex, cell pellets from 1-liter culture were thawed at room temperature and suspended in the buffer containing 20 mM HEPES pH 7.3, 50 mM NaCl, 5 mM CaCl_2_, 5 mM MgCl_2_ with 100× concentrated EDTA-free protease inhibitor cocktail (Bimake). The suspensions were treated with French Press and added with 5 μM His8-SUMOstar-C3a (homemade) and 25 mU/ml apyrase (Sigma), followed by incubation for 1.5 hours at room temperature. After incubation, the complex was extracted from the membrane with 0.5% (w/v) lauryl maltose neopentylglycol (LMNG, Anatrace) and 0.1% (w/v) cholesteryl hemisuccinate (CHS, Anatrace) for 3 h at 4°C. The supernatant was further isolated by centrifugation at 100,000 g for 45 min and then incubated with pre-equilibrated Nickel-NTA resin (20 mM HEPES pH 7.3, 100 mM NaCl) overnight at 4 °C. The nickel resin was loaded onto a gravity column manually. The resin was firstly washed with 15 column volumes of 20 mM HEPES, pH 7.3, 100 mM NaCl, 30 mM imidazole, 1 μM His8-SUMOstar-C3a, 0.01% LMNG (w/v), 0.002% CHS (w/v) and 0.1% digitonin (w/v, Anatrace), and then eluted with the same buffer with 300 mM imidazole in addition for 6 column volumes. The eluted protein was further incubated with M1 anti-FLAG affinity resin (Smart-Lifesciences) with 2 mg scFv16 (homemade) added for 2 h at 4 °C. After incubation, the M1 anti-FLAG affinity resin was washed with 10 column volumes of 20 mM HEPES, pH 7.3, 100 mM NaCl, 1 μM His8-SUMOstar-C3a, 0.01% (w/v) LMNG, 0.01% GDN and 0.002% (w/v) CHS, 0.05% digitonin (w/v) and then eluted with 5 column volumes of the same buffer plus 0.2 mg/mL FLAG peptide. The eluted protein was concentrated to 500μL with a 100 kDa molecular weight cut-off concentrator (ThermoFisher). Concentrated C3a-C3aR-G_i_ complex was loaded onto a Superdex 200 increase 10/300 GL column (GE Healthcare) with running buffer containing 20mM HEPES pH 7.3, 100mM NaCl, 0.00075% LMNG, 0.00025% GDN, 0.0002% CHS,0.05% digitonin. The fractions for monomeric complex were collected, evaluated by SDS-PAGE and concentrated to 11.4 mg/mL for cryo-EM experiments. For apo-C3a-G_i_ complex, the same steps were performed without addition of His6-SUMOstar-C3a, the final sample was concentrated to 16 mg/mL for cryo-EM experiments.

For the purification of C5a-C5aR1-G_i_ complex, no additional C5a ligand was added during the purification of C5a-C5aR1-G_i_ complex. The purify methods were similar to the method of apo-C3a-G_i_ complex. And the final monomeric C5a-C5aR-G_i_ sample was concentrated to 12 mg/mL for cryo-EM experiments.

### Cryo-EM grid preparation and data collection

For the cryo-EM grids preparation, 3 μL of the purified protein complex were applied individually onto EM grids and blotted in a Vitrobot chamber (FEI Vitrobot Mark IV) of 100% humidity at 4 °C. For apo-C3aR-G_i_ and C5a-C5aR1-G_i_ complexes, holey carbon grids (Quantifoil, 300 mesh Au R0.6/1) glow-discharged for 50 seconds were used for EM grids preparation. For C3a-C3aR-G_i_ complex, we used the Au grids (Quantifoil, 300 mesh Au R1.2/1.3) pretreated by glow-discharging and cysteine. Briefly, the Au grids were glow-discharged for 10 seconds. Subsequently, the grids were then transferred to 0.5M cysteine solution for incubation at 30 minutes and washed by ddH2O and anhydrous ethanol, respectively. The treated Au grids were placed in room temperature for 3 minutes before EM grid preparation. The samples were blotted for 2 s and vitrified by plunging into liquid ethane. Grids were stored in liquid nitrogen for condition screening and further data collection.

For the apo-C3aR-G_i_ complex, automatic cryo-EM movie stacks were collected by a Titan Krios G4 at 300KV accelerating voltage equipped with Falcon4 detector in Advanced Center for Electron Microscopy, Shanghai Institute of Materia Medica, Chinese Academy of Sciences (Shanghai, China). The movie stacks were collected automatically with a nominal magnification of 75,000× in counting mode at a pixel size of 0.52 Å. Each movie stack was dose-fractionated in 160 frames with 50 total doses (e/ Å^2^) and collected within a defocus ranging from −0.5 to −2.0 μm. A total of 4,240 movies for the dataset of apo-C3aR-G_i_ complex were collected. Data collection was performed using EPU with one exposure per hole on the grid squares.

For the C3a-C3aR-G_i_ and C5a-C5aR1-G_i_ complex, automatic cryo-EM movie stacks were collected on an FEI Titan Krios microscope operated at 300kV in Advanced Center for Electron Microscopy, Shanghai Institute of Materia Medica, Chinese Academy of Sciences (Shanghai, China). The microscope was equipped with a Gatan Quantum energy filter. The movie stacks were collected automatically using a Gatan K3 direct electron detector with a nominal magnification of 105,000× in super-resolution counting mode at a pixel size of 0.412 Å. The energy filter was operated with a slit width of 20 eV. Each movie stack was dose-fractionated in 36 frames with a dose of 1.39 electrons per frame and collected within a defocus ranging from −0.8 to −1.8 μm. The total exposure time was 2.35 s. A total of 6,762 movies was collected for C3a-C3aR-G_i_ complex. A total of 5,006 movies was collected for C5a-C5aR1-G_i_ complex. Data collection was performed using EPU with one exposure per hole on the grid squares.

### Data processing and 3D reconstruction

Movie stacks were subjected to beam-induced motion correction using MotionCor 2.1 ^64^. Contrast transfer function (CTF) parameters for each non-dose-weighted micrograph were determined by Ctffind4 ^65^. Automated particle selection and data processing were performed using RELION-3.1 beta2 ^66^ or RELION-4.0 beta ^67^.

For the datasets of apo-C3aR-G_i_ complex, the movie stack was aligned, dose weighted, and binned by 2 to 1.6 Å per pixel. The micrographs with resolution worse than 4.0 Å and micrographs imaged within the carbon area of grid squares were abandoned, producing 4,221 micrographs to do further data processing. Template-based particle selection yielded 4,404,753 particles which were subjected to reference-free 2D classifications to discard bad particles. The map of WKYMVm-FPR2-G_i_-scFv16 complex (EMDB: EMD-20126) ^68^ low-pass filtered to 60 Å was used as a reference model for two rounds of maximum-likelihood-based 3D classifications and produced 275,847 particles. These particles were subsequently subjected to 3D refinement, CTF refinement, Bayesian polishing and DeepEMhancer ^69^, which generated a density map with an indicated global resolution of 3.2 Å at a Fourier shell correlation of 0.143.

For the datasets of C3a-C3aR-G_i_ and C5a-C5aR1-G_i_ complex, the movie stack was aligned, dose weighted and binned by 2 to 0.84 Å per pixel. The micrographs with resolution worse than 4.0 Å and micrographs imaged within the carbon area of grid squares were abandoned, producing 4,847 micrographs for C5a-C5aR1-G_i_ complex to do further data processing. For the C3a-C3aR-G_i_ complex, template-based particle selection yielded 5,864,752 particles which were subjected to reference-free 2D classifications to discard bad particles. The map of apo-C3aR-G_i_ low-pass filtered to 40 Å was used as a reference model for five rounds of maximum-likelihood-based 3D classifications. Further 3D classification focusing on the receptor produced 246,827 particles. These particles were subsequently subjected to 3D refinement, CTF refinement, Bayesian polishing and DeepEMhancer which generated a density map with an indicated global resolution of 2.9 Å at a Fourier shell correlation of 0.143. For C5a-C5aR1-G_i_ complex, template-based particle selection yielded 4,236,350 particles which were subjected to reference-free 2D classifications to discard bad particles. The map of C3a-C3aR-G_i_ complex low-pass filtered to 60 Å was used as a reference model for three rounds of maximum-likelihood-based 3D classifications and 3D classification focused on receptor resulting in final subset with 406,559 particles. These particles were subsequently subjected to 3D refinement with scFv16 masked, CTF refinement, Bayesian polishing and DeepEMhancer, which generated a density map with an indicated global resolution of 3.0 Å at a Fourier shell correlation of 0.143.

### Model building, structure refinement, and figure preparation

The G_i_ structure of μOR-TRV130-G_i_ complex (PDB: 8EFB) was used for the model building of this study. The starting models of C3aR and C5aR1 was generated by Alphafold2 ^70^. The start model of C3a and C5a was referenced of previous crystal structures (C3a PDB: 4HW5) (C5a PDB: 5B4P). The structural model was firstly docked as a rigid body into the cryo-EM density maps using UCSF Chimera ^71^. Models were then manually rebuilt and/or adjusted in COOT ^72^. Real space and Rosetta refinements were performed using Phenix ^73^. The model statistics were validated using MolProbity ^74^. Structural figures were prepared in Chimera and PyMOL (https://pymol.org/2/). The final refinement statistics are provided in table S1. The maximum distance cutoffs for polar hydrogen-bond interactions and hydrophobic interactions were set at 3.5 Å and 4.5 Å, respectively.

### GloSensor cAMP assay

The N-terminus of full-length C3aR and C5aR1 was fused with HA signal peptide and FLAG epitope. The constructs were cloned into pcDNA3.0 vector for HEK293T system. Before transfection, HEK293T cells were plated onto 6-well plate with density of 2×10^5^ cells/mL. After 16 hours, cells were transfected with 1.5 μg receptor and 1 μg GloSensor-22F (Promega). After 24 hours, transfected cells were digested and transferred onto 96-well plate with 50 μL suspension with density of 5×10^5^ cells/ml. After another 16 hours, cells were starved by 50 μL Hank’s balanced salt solution for 30 min and then incubated in 50 μL CO_2_-independent media containing 2% GloSensor cAMP Reagent (Promega) for 1 hour. After incubation, 5.5 μL test ligands with various concentrations were added and incubated for 10 min at room temperature. Then 5.5 μL Forskolin were added to the cells in the final concentration of 1 μM. All luminescence signals are tested by EnVision multi-plate reader according to the manufacturer’s instructions. All data were analyzed using Prism 9 (GraphPad) and presented as means ± S.E.M. from at least three independent experiments in technical duplicates or triplicates. The top value was normalized to 0% and the bottom value was normalized to 100% for the final presentation. Non-linear curve fit was performed using a three-parameter logistic equation [log (agonist vs response)]. The significance was determined with two-side, one-way ANOVA followed by Fisher’s LSD test compared with WT. *P<0.05; **P<0.01 and ***P<0.001 were considered as statistically significant.

## Acknowledgements

The cryo-EM data were collected at the Advanced Center for Electron Microscopy, Shanghai Institute of Materia Medica (SIMM). We sincerely thank all the staffs at the institution for their assistance in cryo-EM data collection. This work was partially supported by grants from the Ministry of Science and Technology (China) grants (2018YFA0507002 to H.E.X.); the National Natural Science Foundation of China (82121005 to H.E.X.), the Shanghai Municipal Science and Technology Major Project (2019SHZDZX02 to H.E.X.); the CAS Strategic Priority Research Program (XDB08020303 to H.E.X.); the Special Research Assistant Project of Chinese Academy of Sciences (to Y.W.Z.).

## Authors Contributions

Y.W. designed the expression constructs of C3aR and C5aR1, performed data acquisition and structure determination of C5a-C5aR1-G_i_-scFv16, performed cAMP assays, and participated in figure preparation and manuscript editing. W.Y.L. optimized the purification conditions of protein complexes and prepared protein samples of apo-C3aR-G_i_-scFv16, C3a-C3aR-G_i_-scFv16, and C5a-C5aR-G_i_ complexes for cryo-EM grid making and data collection and participated in method preparation. Y.W.Z. performed data acquisition and structure determination of apo and C3a-bound C3aR-G_i_-scFv16 complex. Y.W.X., Q.N.Y. and Y.W.Z. built the models and refined the structures. X.H.H. performed the molecular dynamic simulation and homology modeling work. P.L., W.J.F., J.P.Z. and X.Y.Z. assisted in cloning construction and protein sample preparation. X.C. supervised X.H.H. in the computational analysis. Y.J. supervised Y.W. and W.Y.L. Y.W.Z. and H.E.X. conceived and supervised the project and wrote the manuscript. Y.W.Z. prepared the draft of the manuscript with the inputs from Y.W. and W.Y.L.

---

**Extended Data Fig. 1.**
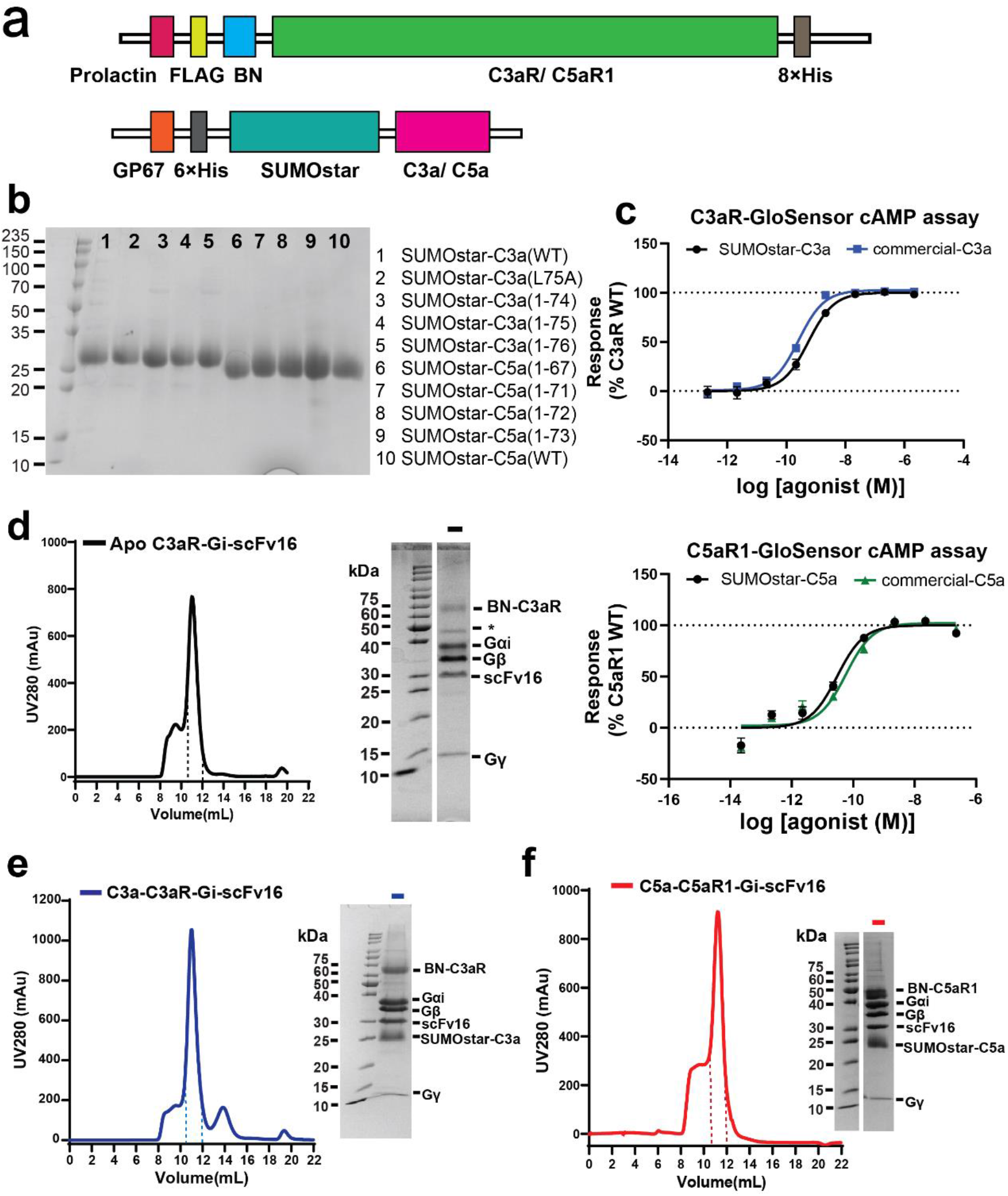
Biochemical results of complement system in this study. **a,** Schematic representation of the C3aR/C5aR1 and C3a/C5a constructs. **b,** SDS-PAGE analysis of the recombinant C3a/C5a mutants and truncations. **c,** Comparisons of the capabilities of homemade C3a and C5a in C3aR and C5aR1 activation using the commercially available C3a (upper panel) and C5a (lower panel) as reference ligands. **d, e, f,** Size exclusion chromatography profiles (left) and SDS-PAGE analysis (right) of the apo-C3aR-G_i_ complex **(d)**, C3a-C3aR-G_i_ complex **(e)** and C5a-C5aR1-G_i_ complex **(f)**.

**Extended Fig. 2.**
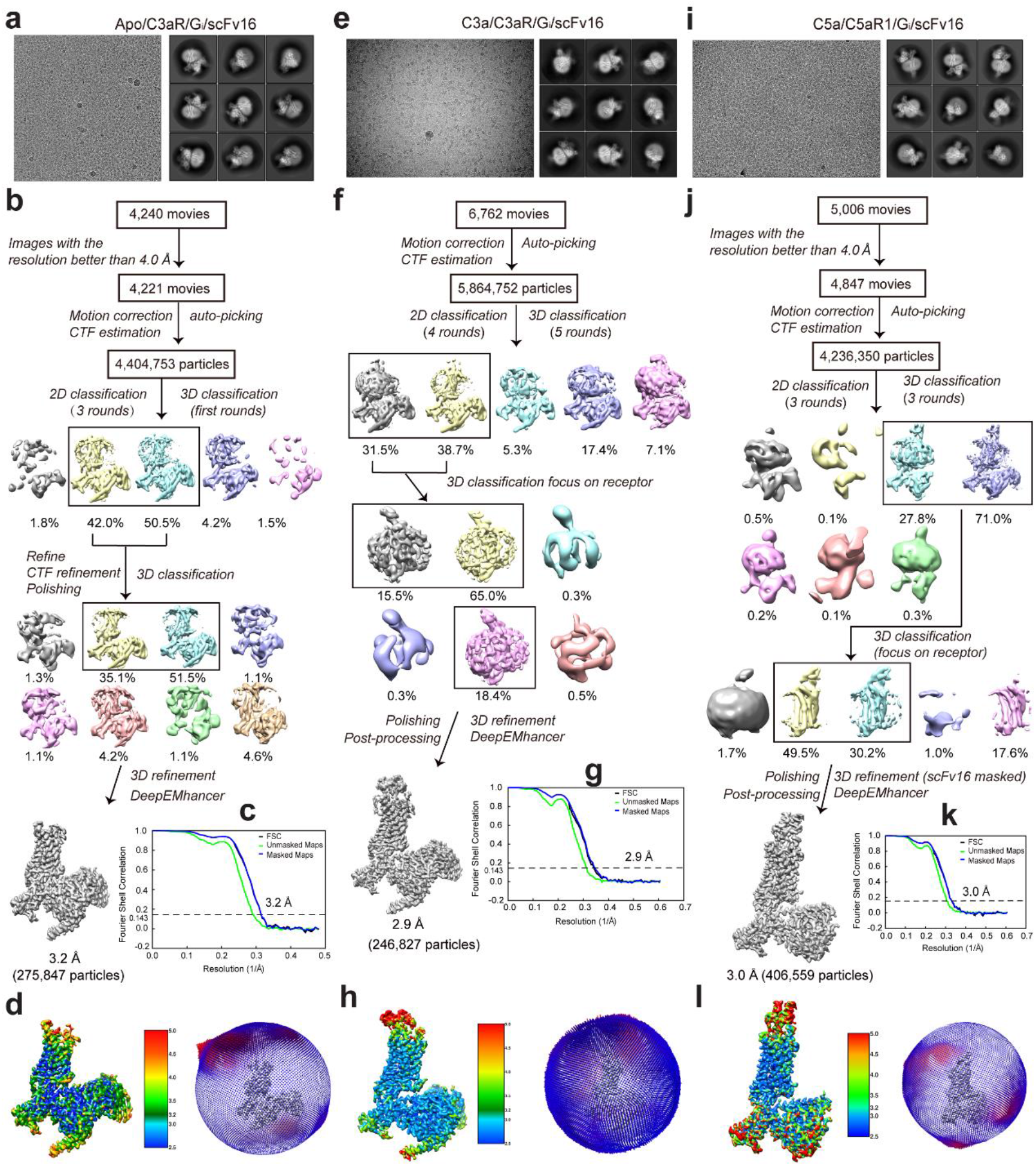
Structure determination of the apo/C3a-C3aR-G_i_, and C5a-C5aR1-G_i_ complex. **a,** Representative cryo-EM raw image and 2D classification averages of the apo-C3aR-G_i_ complex. **b,** Cryo-EM data processing flowchart of the apo-C3aR-G_i_ complex. **c,** The Fourier shell correlation (FSC) curves of the apo-C3aR-G_i_ complex. The global resolution of the final processed density map estimated at the FSC=0.143 is 3.2 Å. **d,** Local resolution and angle distribution map of the apo-C3aR-G_i_ complex. The density map is shown at 0.08 threshold. **e,** Representative cryo-EM image and 2D classification averages of the C3a-C3aR-G_i_ complex. **f,** Cryo-EM data processing work-flow of the C3a-C3aR-G_i_ complex. **g,** The Fourier shell correlation (FSC) curves of the apo-C3aR-G_i_ complex. The global resolution of the final processed density map estimated at the FSC=0.143 is 2.9 Å. **h,** Local resolution and angle distribution map of the C5a-C5aR1-G_i_ complex. The density map is shown at 0.25 threshold. **i,** Representative cryo-EM image and 2D classification averages of the C5a-C5aR1-G_i_ complex. **j,** Cryo-EM data processing flowcharts of the C5a-C5aR1-G_i_ complex. **k,** The Fourier shell correlation (FSC) curves of the C5a-C5aR1-G_i_ complex. The global resolution of the final processed density map estimated at the FSC=0.143 is 3.0 Å. **l,** Local resolution and angle distribution map of the C5a-C5aR1-G_i_ complex. The density map is shown at 0.11 threshold.

**Extended Fig. 3.**
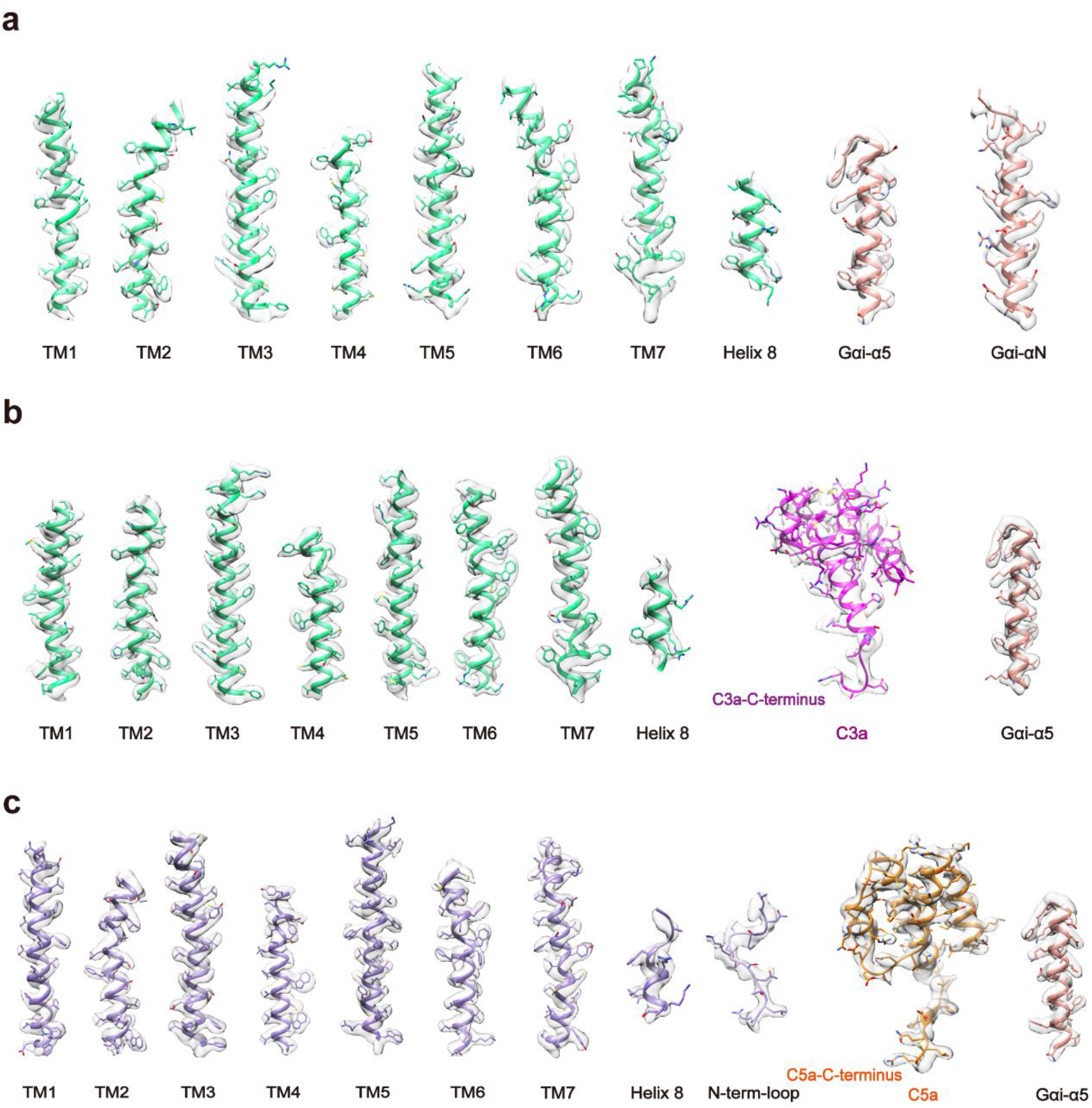
Local electron densities of C3aR-G_i_ and C5aR1-G_i_ complexes. **a, b, c,** EM density maps of transmembrane helices TM1-TM7 and helix 8 of C3aR or C5aR1, αN or α5 helices of G_i_, and ligands C3a and C5a in the apo-C3aR-G_i_ complex **(a)**, the C3a-C3aR-G_i_ complex **(b)**, and the C5a-C5aR1-G_i_ complex(**c**). The density maps were shown at the thresholds of 0.08, 0.15 and 0.08 for apo-C3aR-G_i_ complex, the C3a-C3aR-G_i_ complex, and the C5a-C5aR1-G_i_ complex, respectively.

**Extended Fig. 4.**
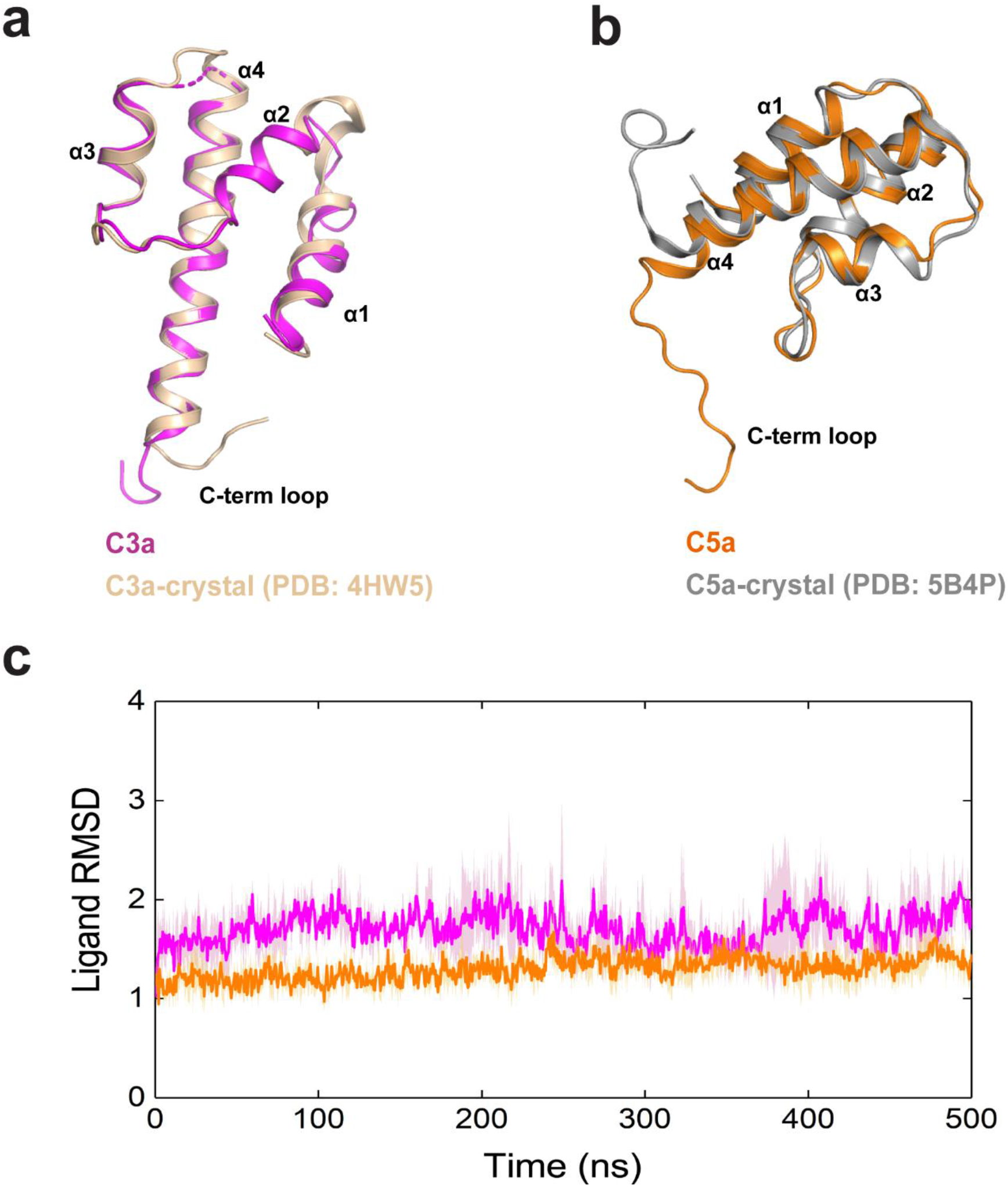
Molecular dynamic simulations of C3a and C5a binding poses. **a,** Superposition of C3a structure determined by cryo-EM in this study and crystal structure of C3a (PDB: 4HW5). **b,** Superposition of C5a structure determined by cryo-EM in this study and crystal structure of C5a (PDB: 5B4P). **c,** Molecular dynamics simulations of C3a and C5a bound to C3aR and C5aR1, respectively.

**Extended Fig. 5.**
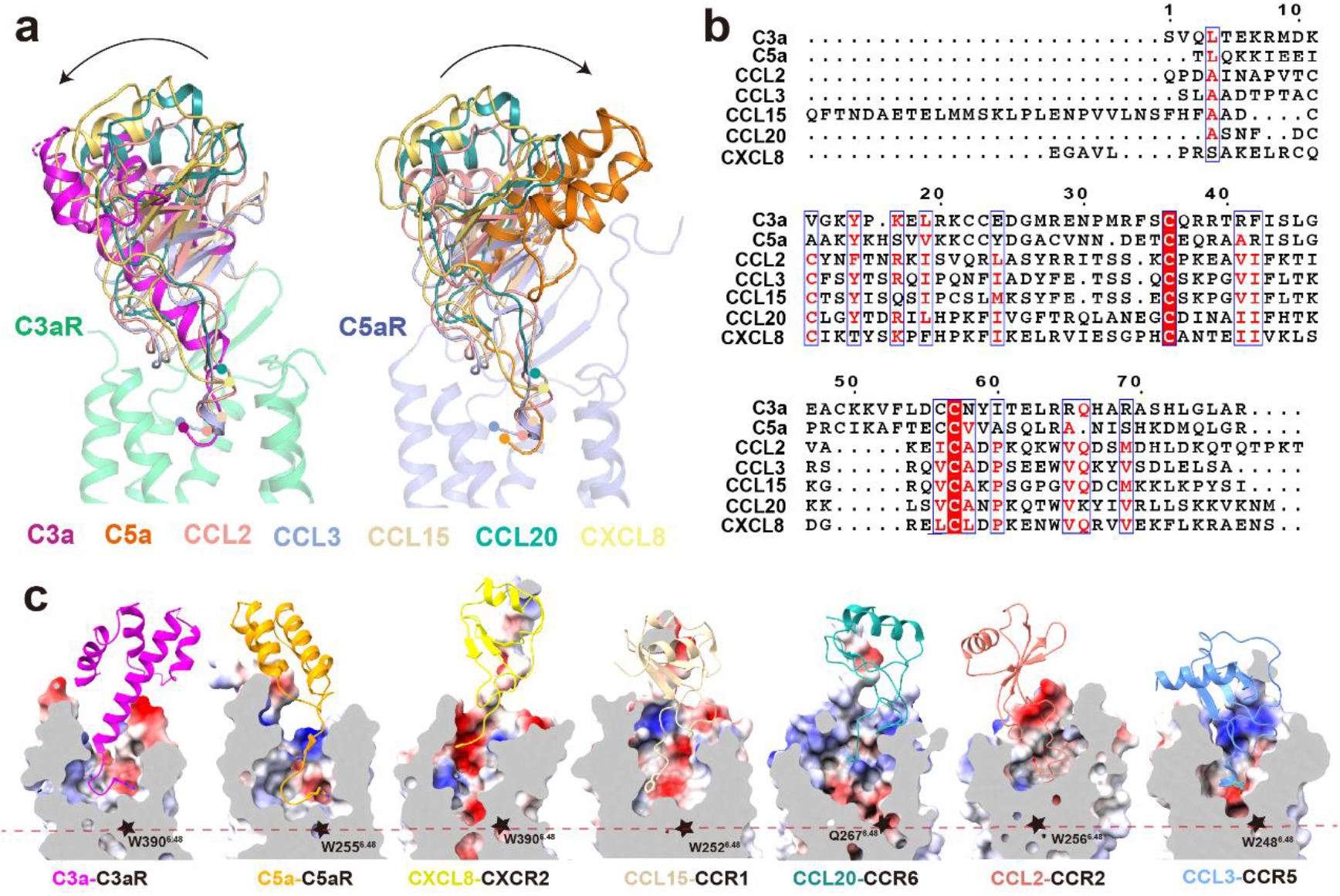
Distinct binding modes of C3a and C5a from chemokines. **a,** Structural superposition of C3a bound C3aR and C5a bound C5aR1 with chemokine receptors, only chemokines and C3aR/C5aR1 were shown in cartoon. CCL2-CCR2 (PDB: 7XA3), CCL3-CCR5 (PDB: 7F1Q), CCL15-CCR1 (PDB: 7VL9), CCL20-CCR6 (PDB: 6WWZ), CXCL8-CXCR2 (PDB: 6LFO). **b,** Amino acid sequence alignment of C3a/C5a with chemokines CCL2, CCL3, CCL15, CCL20 and CXCL8. **c,** Cross-section of ligand binding pockets of anaphylatoxin receptors and chemokine receptors. Stars indicate the conserved tryptophan residue at position 6.48.

**Extended Fig. 6.**
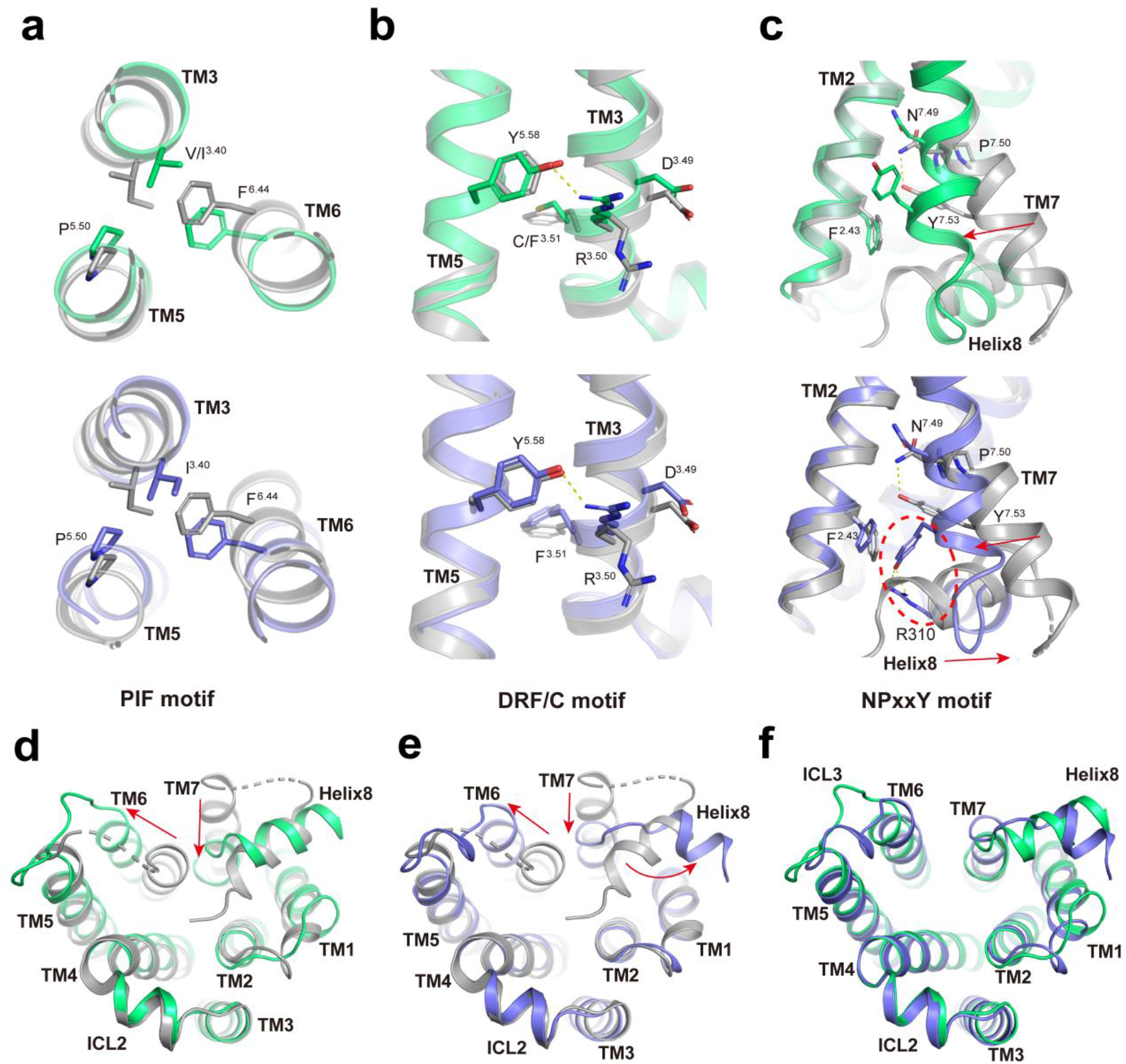
Conformational changes of C3aR and C5aR1 activation. **a, b, c,** Conformational changes upon activation of C5aR1 induced by C5a, including rearrangement of PIF motif **(a)**, alteration of DRF motif **(b)** and NPxxY motif **(c)**. **d, e, f,** Conformational changes upon activation of C3aR induced by C3a, including rearrangement of PIF motif **(d)**, alteration of DRC motif **(e)** and NPxxY motif **(f)**. **g, h, i,** Intracellular region conformation changes when super-positioned C3a bound C3aR with inactive C5aR1 **(g)**, C5a bound C5aR1 with inactive C5aR1 **(h)** and C3a bound C3aR with C5a bound C5aR1 **(i)**.

**Extended Fig. 7.**
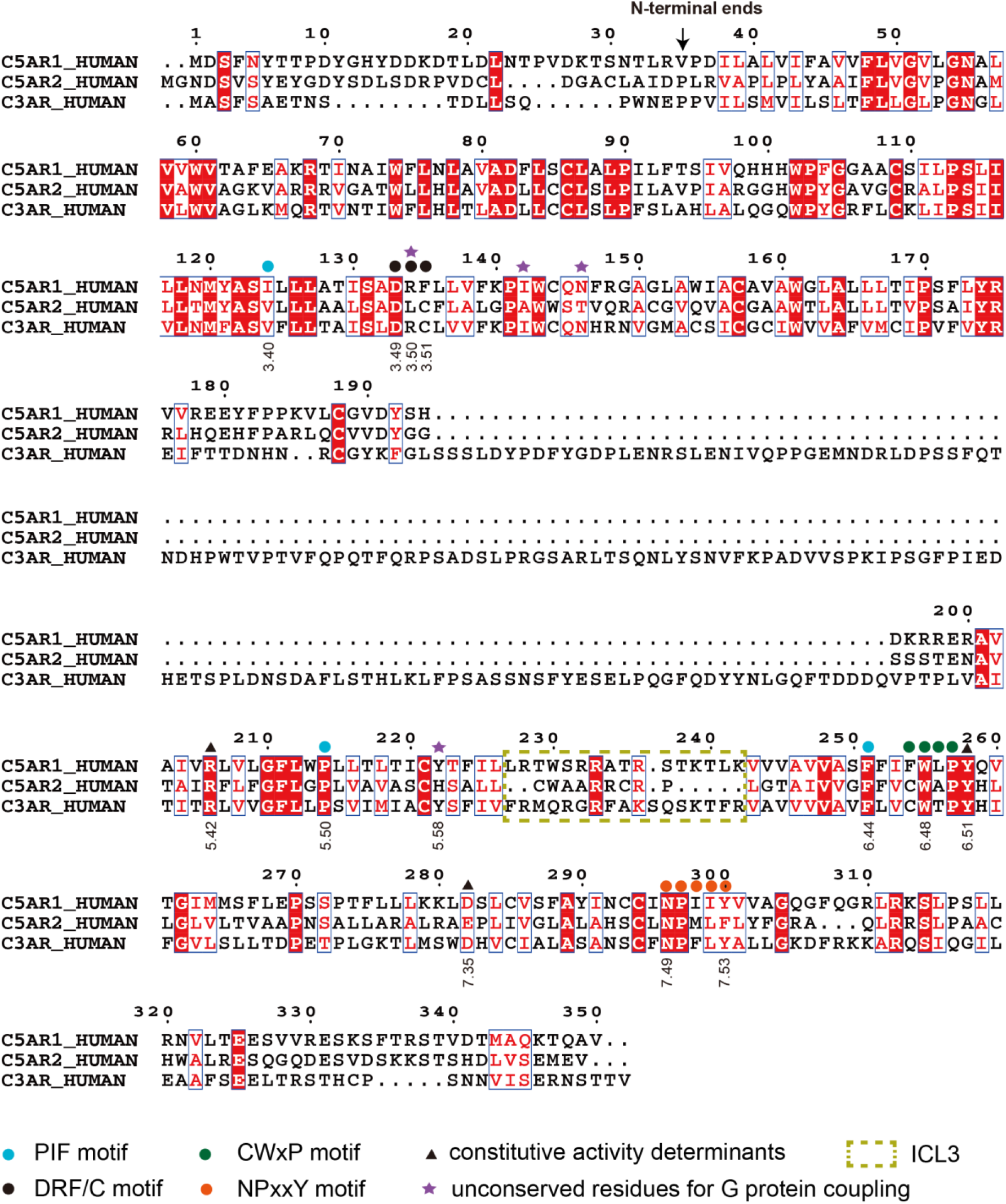
Sequence alignment of the anaphylatoxin receptors. The sequences shown are those for human C3aR, C5aR1 and C5aR2 which was created using Clustalw and ESPript 3.0 servers.

**Extended Fig. 8.**
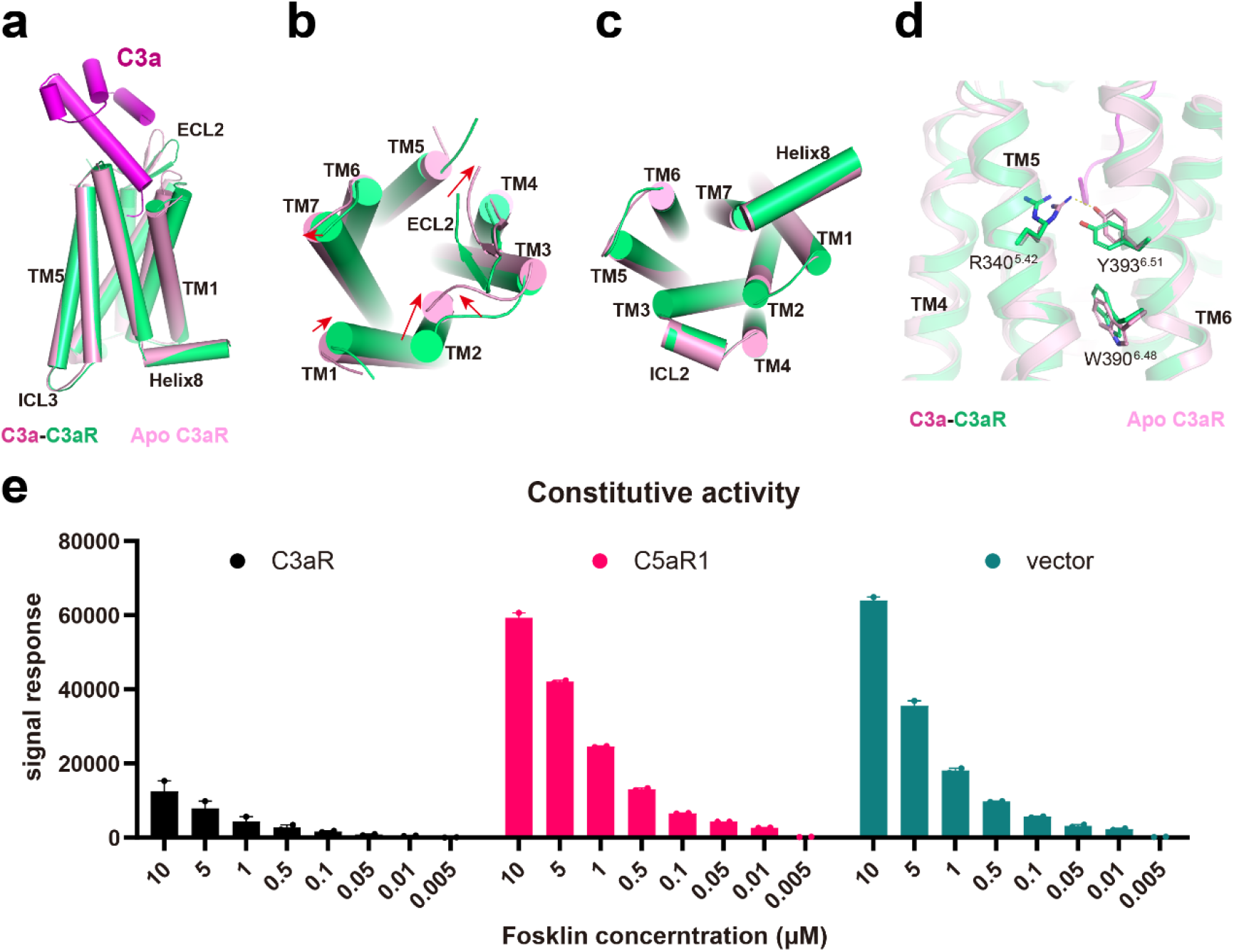
Constitutive activity determinants of C3aR. **a, b, c,** Structural superposition of the C3a-C3aR with the apo-C3aR in orthogonal view **(a)**, extracellular view **(b)** and intracellular view **(c)**. Helixes are shown as rods. **d,** Constitutive determinant residues of C3aR. In apo-C3aR, residue R340^5.42^ forms direct hydrogen bond with Y393^6.51^ and keeps it in active conformation. **e,** Histogram of constitutive activity of C3aR and C5aR1, controlled as pcDNA3.0 vector. Cells were treated with decreasing dose of Fosklin. It can be seen that C3aR has high basal activity whereas C5aR1 has no basal activities and behaves like the control.

**Extended Fig. 9.**
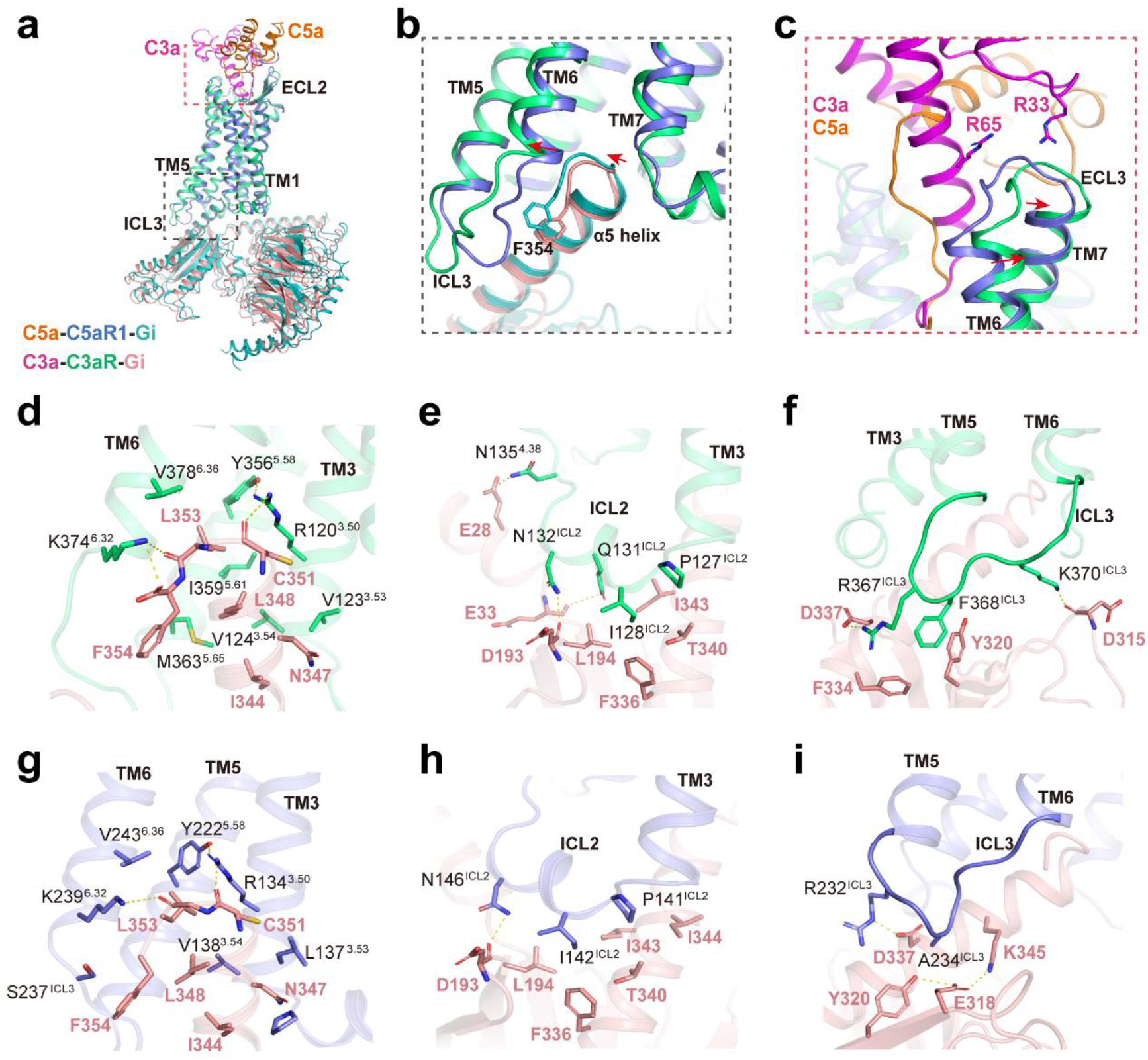
G_i_ coupling of C3aR and C5aR1. **a,** Overall structural superposition of the C3a-C3aR-G_i_ complex with the C5a-C5aR1-G_i_ complex. **b,** Subtle differences of α5 helix of G_αi_ subunit inserted into C3aR and C5aR1. **c,** C3a/C5a induced the extracellular region movement between C3aR and C5aR1. **d, e, f,** Interactions between C3aR and G_αi_ subunit, **(d)** intracellular cavity of C3aR with α5 helix of G_αi_ subunit, **(e)** ICL2 of C3aR with G_αi_ subunit, **(f)** ICL3 of C3aR with G_αi_ subunit. **g, h, i,** Interactions between C5aR1 and G_αi_ subunit, **(g)** intracellular cavity of C5aR1 with α5 helix of G_αi_ subunit, **(h)** ICL2 of C5aR1 with G_αi_ subunit, **(i)** ICL3 of C5aR1 with G_αi_ subunit.

